# High intraspecies allelic diversity in *Arabidopsis* NLR immune receptors is associated with distinct genomic and epigenomic features

**DOI:** 10.1101/2023.01.12.523861

**Authors:** Chandler A. Sutherland, Daniil M. Prigozhin, J. Grey Monroe, Ksenia V. Krasileva

## Abstract

Plants rely on Nucleotide-binding, Leucine-rich repeat Receptors (NLRs) for pathogen recognition. Highly variable NLRs (hvNLRs) show remarkable intraspecies diversity, while their low variability paralogs (non-hvNLRs) are conserved between ecotypes. At a population level, hvNLRs provide new pathogen recognition specificities, but the association between allelic diversity and genomic and epigenomic features has not been established. Our investigation of NLRs in *Arabidopsis* Col-0 has revealed that hvNLRs show higher expression, less gene body cytosine methylation, and closer proximity to transposable elements than non-hvNLRs. hvNLRs show elevated synonymous and nonsynonymous nucleotide diversity and are in chromatin states associated with an increased probability of mutation. Diversifying selection maintains variability at a subset of codons of hvNLRs, while purifying selection maintains conservation at non-hvNLRs. How these features are established and maintained, and whether they contribute to the observed diversity of hvNLRs is key to understanding the evolution of plant innate immune receptors.

## Introduction

Plants, lacking the adaptive immune systems of vertebrates, use germline-encoded innate immune receptors to defend against rapidly evolving pathogens. Despite their inability to create antibodies through somatic hypermutation and recombination, plants are protected against pathogens due to population-level receptor diversity. Nucleotide-binding, Leucine-rich repeat Receptors (NLRs) are the intracellular sensors of the plant immune system, detecting pathogen-secreted, disease-promoting effector proteins. After binding of a pathogen target to the LRR domain, NLRs initiate defense responses through oligomerization of the central nucleotide-binding domain, leading to transcriptional reprogramming, hormone induction, and hypersensitive cell death response (Ngou, Ding and Jones, 2022). Plant NLRs are differentiated into three anciently diverged classes based on their N-terminal domains: Resistance To Powdery Mildew 8-NLR (RNL), Coiled-Coil-NLR (CNL), or Toll/Interleukin-1 Receptor-NLR (TNL) that are responsible for the downstream signaling.

NLRs exhibit remarkable levels of intraspecies allelic diversity (Van de Weyer *et al*., 2019), due to both the genomic processes that generate variation and selection that promotes its maintenance (Karasov, Horton and Bergelson, 2014; Barragan and Weigel, 2021; Märkle, Saur and Stam, 2022). NLRs are organized into clusters more often than other genes, which can asymmetrically drive NLR expansion and diversification through unequal crossing over and gene conversion (Michelmore and Meyers, 1998; Lee and Chae, 2020) as well as accumulation of point mutations (Kuang *et al*., 2004). Point mutations are a major source of within-species NLR diversity, but have been difficult to fully resolve through short-read sequencing approaches. The NLR gene family includes the most polymorphic loci and contains the highest frequency of major effect mutations in the *Arabidopsis* genome (Gan *et al*., 2011). There is evidence for balancing selection maintaining polymorphisms and presence-absence variation at several NLR loci through frequency-dependent selection, spatial and temporal fluctuations in pathogen pressure, and heterozygote advantage (Thrall *et al*., 2012; Karasov *et al*., 2014; MacQueen *et al*., 2019). Diversifying selection has also been observed at NLR loci as an excess of nonsynonymous to synonymous substitutions (Bakker *et al*., 2006). The NLR gene and protein sequences within a species represent a snapshot of the ongoing interplay between mutation and selection, but disentangling their relative contributions remains challenging.

Mutation rates are unlikely to evolve on a gene by gene basis in response to selection given the barrier imposed by genetic drift (Lynch, 2010). However, selection on genic mutation rates is sufficiently strong when acting on mechanisms that couple mutation rate to expression states and epigenomic features, affecting the mutation rates of many genes simultaneously (Martincorena and Luscombe, 2013). The mutation rate of *Arabidopsis* is heterogeneous across the genome, consistent with expected effects of selection on mechanisms linking mutation rates to epigenomic features (Monroe *et al*., 2022; Staunton, Peters and Seoighe, 2023). Several mechanisms have been described, including cytosine methylation which is positively correlated with mutation probability and known to increase the likelihood of spontaneous deamination (Cao *et al*., 2011; Weng *et al*., 2019) while H3K4me1, which is negatively correlated with mutation probability and a target of several DNA repair proteins (Quiroz *et al*., 2022). Description of genomic features associated with diversity in NLRs will help to understand the role of mutation bias in NLR evolution.

Recent advances in enrichment-based long-read sequencing of NLRs (Jupe *et al*., 2013) as well as long-read pan-genomes (Jiao and Schneeberger, 2020) allowed for re-examination of NLR variation within species (Barragan and Weigel, 2021). In *Arabidopsis* datasets, it has been shown that NLRs are enriched in regions of synteny diversity and that NLR repertoires across species could not be easily anchored to a reference genome (Van de Weyer *et al*., 2019). Phylogenetic analysis independent of reference-based assignment of pan-NLRomes from 62 *Arabidopsis thaliana* accessions (Van de Weyer *et al*., 2019) and 54 *Brachypodium distachyon* (Gordon *et al*., 2017) lines allowed for amino acid diversity quantification and delineation of highly variable NLRs (hvNLRs) from their low-variability paralogs (non-hvNLRs) (Prigozhin and Krasileva, 2021). At the population level, hvNLRs show rapid rates of diversification and are hypothesized to act as reservoirs of diversity for recognition of pathogen effectors. Comparison of hv and non-hvNLR gene sets allows for investigation of epigenomic, sequence, and regulatory features (hereafter genomic features) and signatures of selection associated with NLR diversification.

In this paper, we report that hvNLRs show a higher transcription level, less gene body CG methylation, and closer proximity to transposable elements (TEs) than non-hvNLRs. Elevated gene-wide nucleotide diversity, a higher likelihood of mutation, and diversifying selection at a subset of sites promote the high amino acid diversity of hvNLRs, while non-hvNLRs are subject to purifying selection. These findings will serve as a starting point for investigation of the mechanisms that promote and maintain diversity generation in a subset of plant immune receptors.

## Results

Shannon entropy, a measure of variability derived from information theory, provides an unbiased metric of amino acid diversity of a protein within a population (Asti *et al*., 2016; Wang *et al*., 2017). Here, the Shannon entropy is the sum of the frequency of each amino acid times the logarithm of that frequency at each position in a protein sequence alignment, so sites with low variability have low entropy and highly diverse sites have high entropy. When applied to NLRs, this measure is predictive of highly variable effector binding sites (Prigozhin and Krasileva, 2021). Based on the bimodal distribution of Shannon entropy in NLRome, we defined hvNLRs as proteins with 10 or more amino acid positions with Shannon entropy greater than 1.5 bits (**Supplemental Fig. 1**) (Prigozhin and Krasileva, 2021). To examine the relationships between population level diversity and genomic features of a single accession, we plotted Shannon entropy by sequence in Col-0 (**Fig. 1**). As expected, there are functional hvNLRs and non-hvNLRs, with known direct recognition of effectors corresponding to hvNLRs and known indirect recognition to non-hvNLRs. hvNLRs also overlap with dangerous mix genes. Categorizing NLRs into low and high entropy groups allows for binary comparison of features and gene set enrichment analysis to compare NLRs to the rest of the genome.

**Figure 1:**
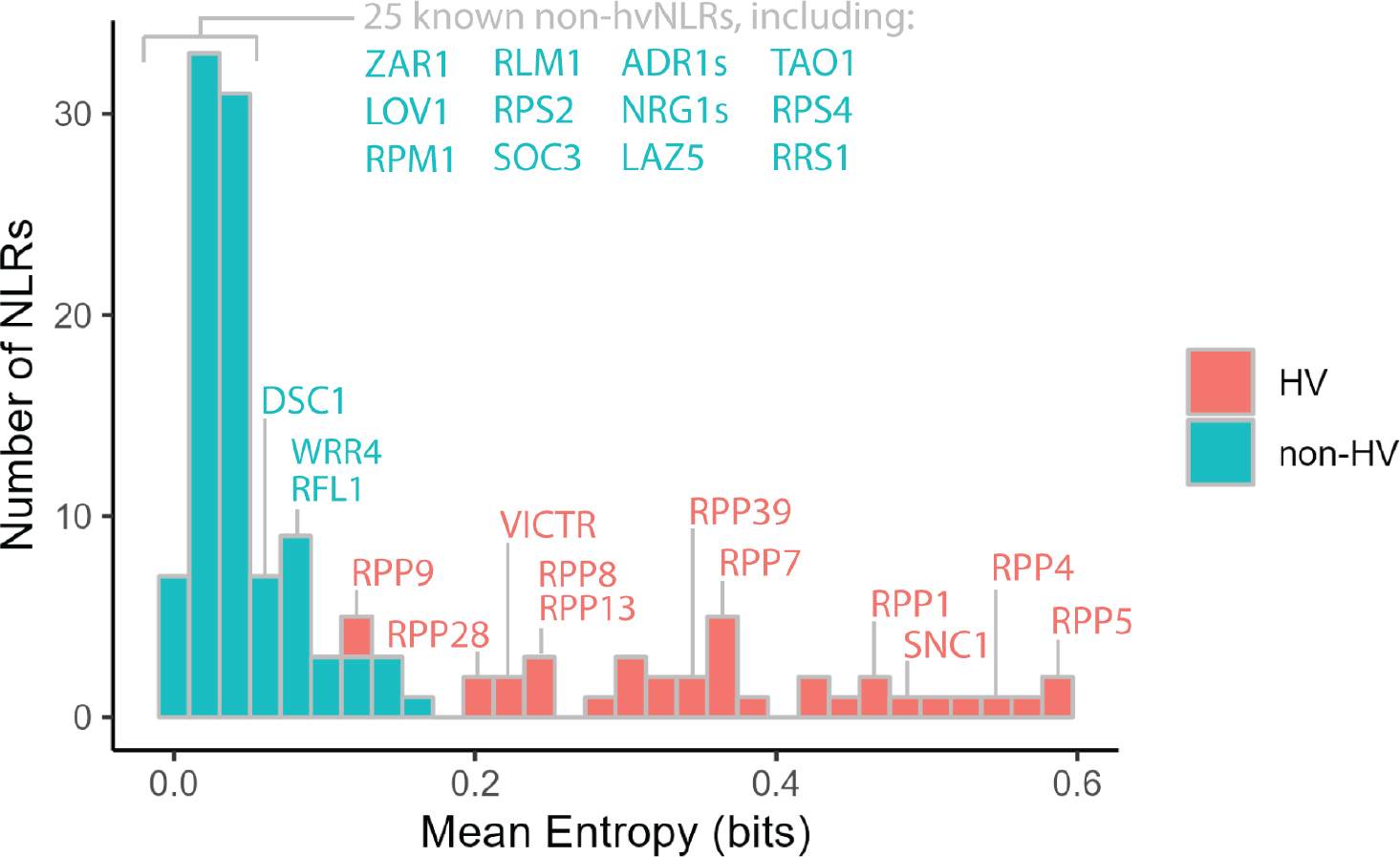
hvNLRs are defined by high amino acid diversity. Distribution of mean per gene Shannon entropy across the *Arabidopsis* NLRome in bits. Described NLRs are annotated.

To compare the expression and methylation status of hv and non-hvNLRs within an individual plant, we examined available paired whole genome bisulfite and RNA sequencing generated from the same rosette leaf (Williams *et al*., 2022). We found that hvNLRs are expressed significantly higher than non-hvNLRs (**Fig. 2A**, unpaired Wilcoxon rank-sum test, p=7.9e-05). When we ranked all protein coding *Arabidopsis* genes based on their expression level, we observed that hvNLRs are enriched in the most expressed genes in each leaf sample (singscore rank-based sample scoring, p < 0.005 for hvNLRs in each biological replicate) (Foroutan *et al*., 2018).

**Figure 2:**
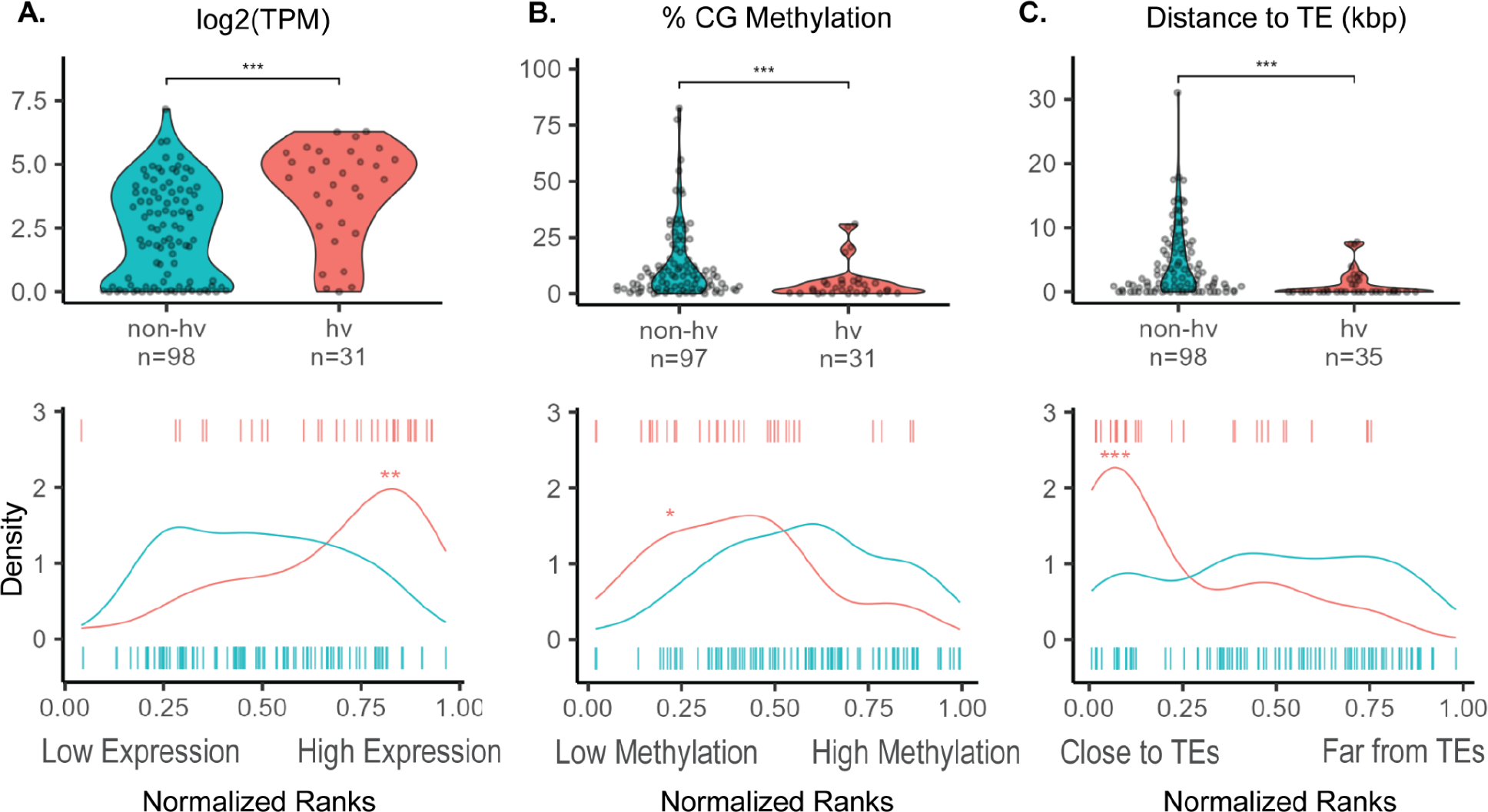
Expression, methylation, and proximity to transposable elements (TEs) distinguish hv and non-hv NLRs. **A:** average gene expression log_2_(Transcripts per Million (TPM)), **B:** average % CG methylation per gene, and **C:** distance to the nearest TE (kbp) with normalized mean percentile rank density plots of hv and non-hvNLRs.* indicates a p-value <0.05 and ≥ 0.01; ** indicates a p-value <.01 and ≥0.001; *** indicates a p-value <0.001.

In addition, hvNLRs have significantly lower gene body CG methylation than non-hvNLRs (**Fig. 2B**, unpaired Wilcoxon rank-sum test, p=4.3e-04), and hvNLRs are enriched in the CG hypomethylated genes across the genome (**Fig. 2B**, permutation test for difference in means, p= 0.003, n=10,000 replicates). Gene set analysis of methylation can be biased due to the uneven distribution of CG sites within each gene (Geeleher *et al*., 2013). We repeated our permutation test to compare hvNLRs to a set of non-NLR genes with similar measured CG sites per gene to correct for this bias. Still, hvNLRs were significantly more hypomethylated than the rest of the genome (p < 0.05 each biological replicate, n=10,000). We noticed two hvNLRs, *RPP4* and *RPP7*, with higher CG methylation than the average for hvNLRs (**Fig. 2B**). Upon further inspection, we also found CHH and CHG context methylation within the gene bodies of *RPP4* and *RPP7* (**Supplemental Fig. 2**), which we rarely observed in other NLRs. Multi-context gene body methylation (CG, CHH, and CHG) is typically used to silence nearby or overlapping transposable elements (Quadrana *et al*., 2016). This indicates that their elevated CG methylation is likely due to multi-context silencing related to a recent TE insertion.

We also found that hvNLRs are much more likely to be near TEs (Fig. 2C, unpaired Wilcoxon rank-sum test, p = 1.7e-06), and hvNLRs are enriched in the genes closest to TEs (permutation test for difference in medians, p=0, n=10,000 replicates). In Col-0, hvNLRs have a median TE distance of 0 kbp, meaning the TEs are within the UTR or intronic sequences, while non-hvNLRs have a median TE distance of 2.07 kbp. Highly variable status of NLRs is predictive of TEs within the genic sequence (Fisher’s exact test, p=3.6e-05). It has been previously observed that TEs are associated with plant immune genes (Kawakatsu *et al*., 2016), but this analysis suggests that the signal is driven by hvNLRs.

NLRs are found in clusters more frequently than other genes (Lee and Chae, 2020). However, highly variable status of NLRs is not dependent on cluster membership (Fisher’s exact test, p=0.18) and hv and non-hvNLRs maintain their distinct expression and TE-association patterns when comparing exclusively clustered hv and non-hvNLRs (**Fig. 3A**, unpaired Wilcoxon rank-sum tests, corrected for multiple hypothesis testing). Expression and TE distance patterns are also independent of the CNL and TNL N-terminal domain clades (**Fig. 3A**). CG methylation, however, is not significantly different between clustered hv and non-hvNLRs and between TNLs (**Fig. 3A**). CG methylation is the weakest association with hv status of the three examined features (**Fig 2B**), and further analysis with more accessions will reveal if cluster or hv status is more predictive of CG methylation. hvNLRs are distributed over the phylogeny of

**Figure 3:**
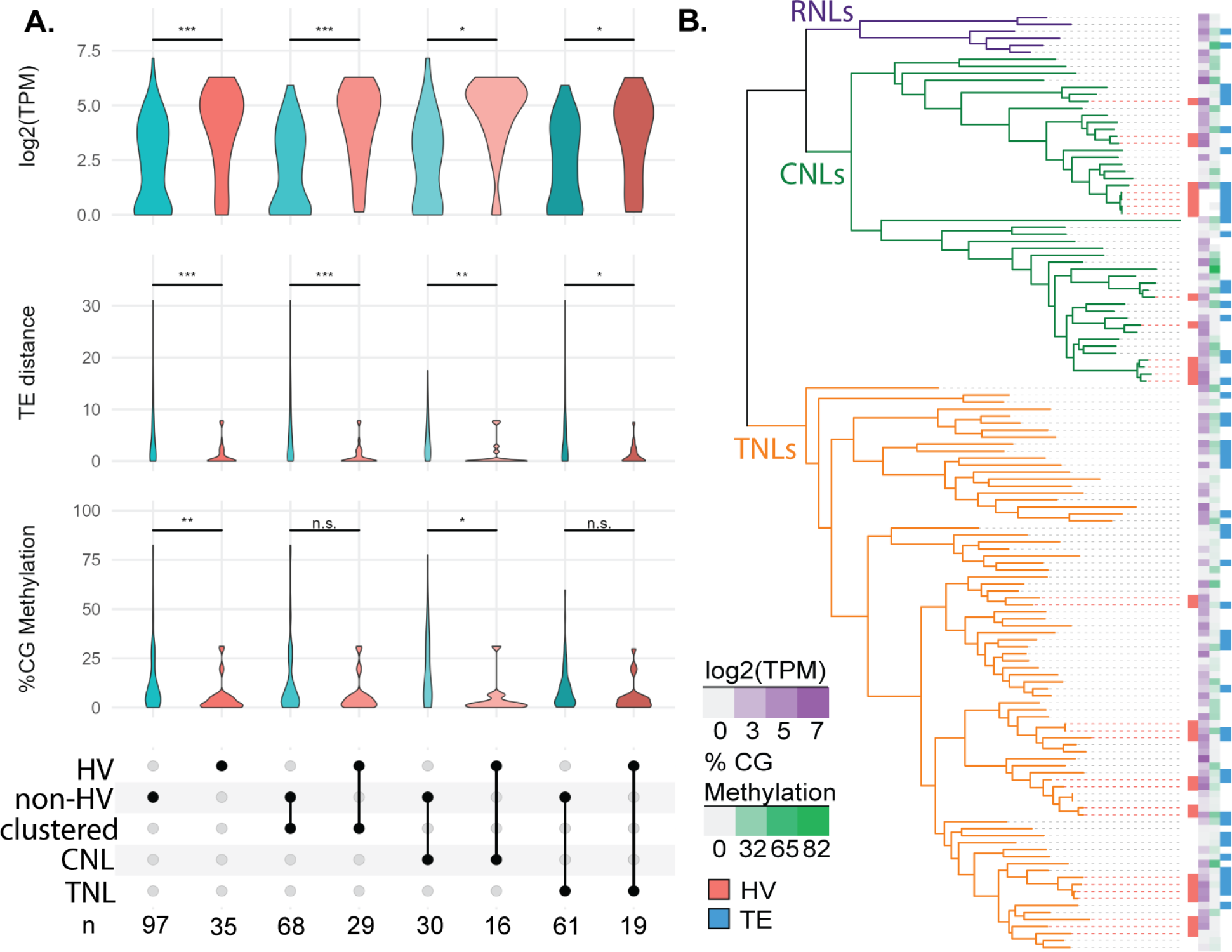
Cluster membership, NLR type, and phylogenetic distance do not account for genomic differences between hv and non-hvNLRs. **A:** Comparison of expression, distance to nearest TEs, and CG gene body methylation of hv and non-hvNLRs by cluster membership and N-term domain type. **B:** Features mapped onto a phylogeny of NLRs in *A. thaliana* Col-0. NLRs without log_2_(TPM) or %CG methylation data were determined to be unmappable (see methods).

NLRs, and maintain distinct genomic features despite close phylogenetic relationships with non-hvNLRs (**Fig. 3B**). Overall, we conclude expression and TE distance cannot be explained by cluster status, phylogenetic proximity, or NLR class.

The high level of amino acid diversity in hvNLRs and associated difference in genomic features might be due to differences in mutational processes and/or selection. In order to investigate the contribution of balancing selection to the observed amino acid diversity at hvNLRs, we calculated Tajima’s D (D) and nucleotide diversity per site (*π*) in each domain and across the gene body of hv and non-hvNLRs. hvNLRs have higher D than non-hvNLRs across the coding sequence and all individual domains (**Fig. 4A**; unpaired Wilcoxon rank-sum test, corrected for multiple comparisons). Reflecting their differences in amino acid diversity, hvNLRs have higher *π* than non-hvNLRs across all domains and the coding sequence (**Fig. 4A**; unpaired Wilcoxon rank-sum test, corrected for multiple comparisons). The difference in *π* and D between the two groups is not driven exclusively by variation in the LRR region, with the highest values reported for the hvNLR NBARC domains.

**Figure 4:**
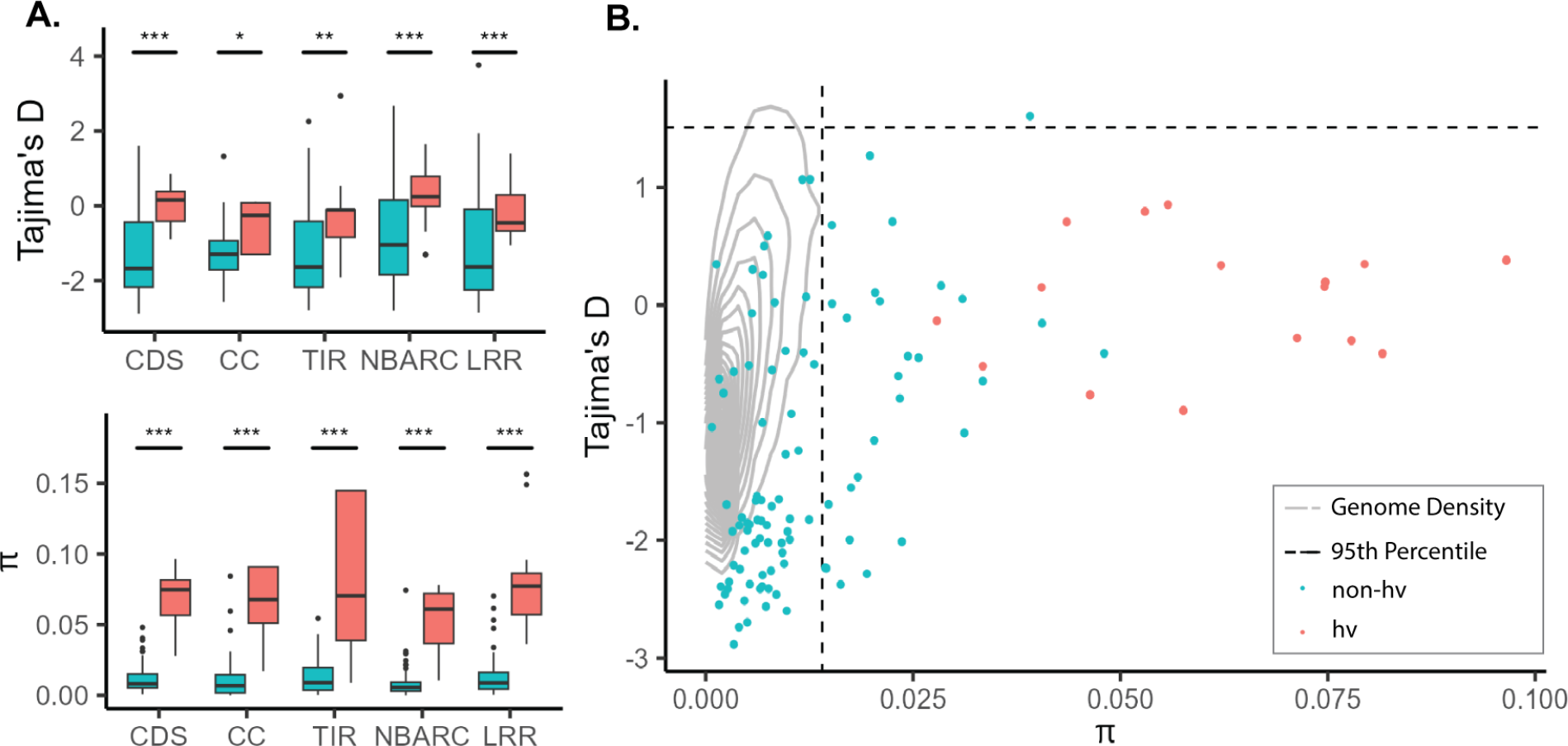
hvNLRs show higher Tajima’s D and nucleotide diversity than non-hvNLRs. **A**. D and *π* calculated across the coding sequence (CDS), coiled-coil (CC), Toll/Interleukin-1 (TIR), nucleotide-binding (NBARC) and leucine rich repeat (LRR) domains. Within each box, horizontal black lines denote median values; boxes range from the 25th to 75th percentile of each group’s distribution of values; whiskers extend no further than 1.5x the interquantile range of the hinge. Data beyond the end of the whiskers are outlying points and are plotted individually. **B**. CDS *π* vs. D. Gray lines represent the kernel density estimation of statistics computed on all coding sequences of *Arabidopsis*. Dashed lines represent the 95th percentile of the empirical distribution.

Due to the demographic history of *Arabidopsis*, the empirical distribution of summary statistics departs from the neutral model (Nordborg et al., 2005; Alonso-Blanco et al., 2016). We calculated the genome-wide values of D and *π* to test for selection, using whole genome SNP information from the accessions used to create the pan-NLRome (Alonso-Blanco *et al*., 2016) (**Supplemental Fig. 3**). Both hv and non-hvNLRs have higher average *π* than the empirical distribution (**Fig. 4B;** permutation test for difference in means, p = 0; p=0, n=10,000 replicates), and there are significantly more NLRs in the top 5% of the empirical distribution than expected by chance (permutation test for number in the top 5%, p=0, n=10,000 replicates). This corroborates previously reported significantly high levels of nucleotide diversity of NLRs. (Bakker *et al*., 2006; Van de Weyer *et al*., 2019) hvNLRs have a higher D and non-hvNLRs have a lower D than the genome average (**Fig. 4B**; permutation test for difference in means, p = 0.0009; p=0, n=10,000 replicates). There are no hvNLRs in either tail of the empirical distribution of D, which is not significantly different from the 0.43 expected by chance. There are, however, an excess of non-hvNLRs in the bottom 5% of the distribution of D (permutation test for number in the bottom 5%, p=0, n=10,000 replicates), indicating that purifying selection may be reducing diversity at non-hvNLRs. Defining individual genes under balancing selection to be the top 5% of the empirical distribution of *π* and D values (Bakker *et al*., 2006; Gladieux *et al*., 2022), we identified one non-hvNLR under balancing selection, *AT5G47260* **(Fig. 4B)**. However, one gene is not significantly different from the number of NLRs expected to be in the top 5% of both distributions by random chance.

To further investigate the nature of the high nucleotide diversity of NLRs, we compared nucleotide diversity at synonymous and non-synonymous sites (*π*_S_;*π*_N_). hvNLRs have greater *π*_S_ and *π*_N_ than non-hvNLRs (**Fig 5A**; unpaired wilcoxon rank sum test, p=5.6e-13, p=1.2e-15). However, the ratio of non-synonymous to synonymous nucleotide diversity (*π*_N_/*π*_S_), an intraspecies measurement of selection, is not significantly different between the two groups, indicating possible role of different mutational processes (**Fig 5B**; unpaired wilcoxon rank sum test, p=0.24). Average *π*_N_/*π*_S_ is < 1 for both groups across the gene and in the LRR region, indicating purifying selection as an excess of synonymous polymorphisms relative to non-synonymous polymorphisms (**Fig. 5B**; **Supplementary Fig. 4**).

**Figure 5:**
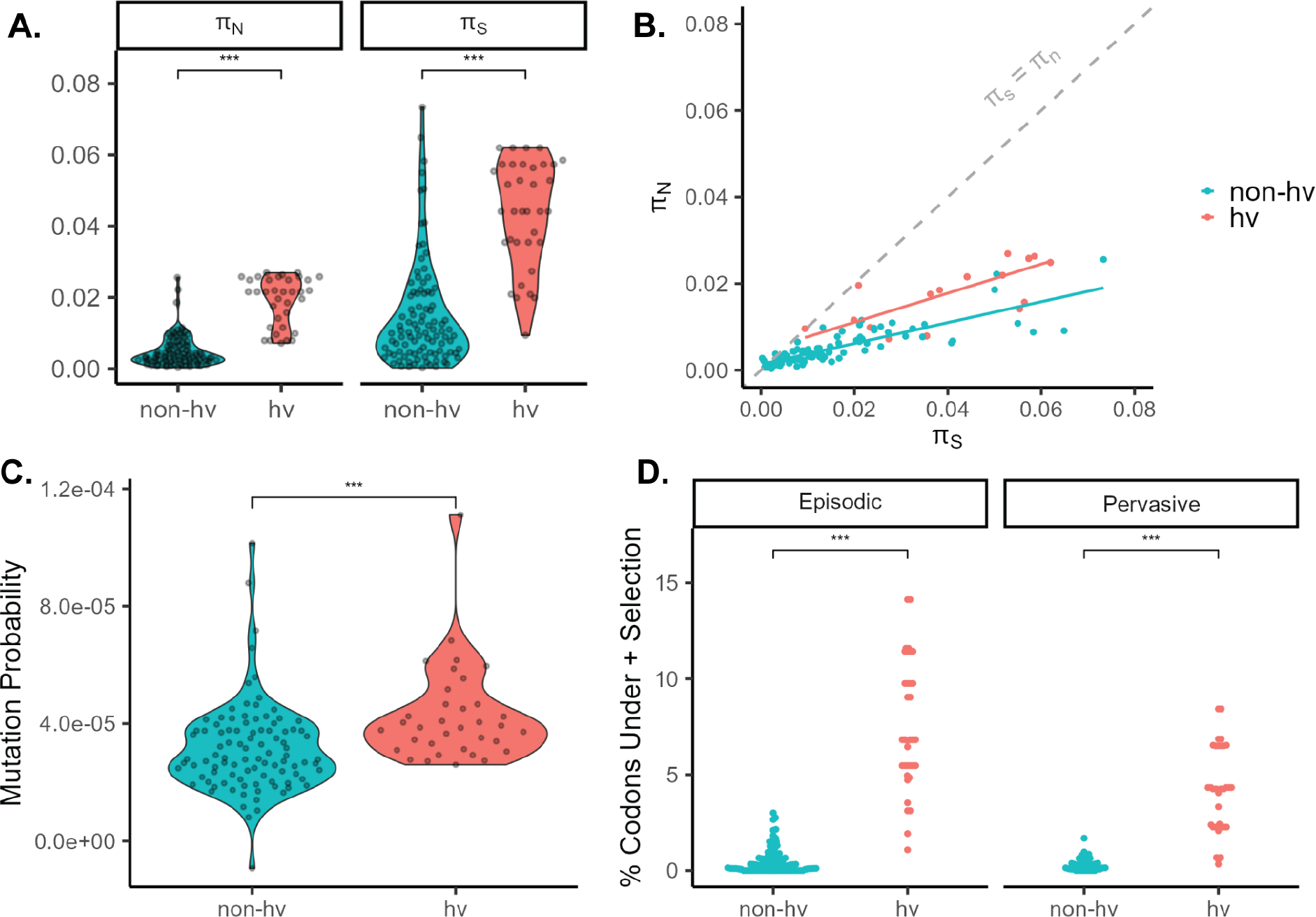
hvNLR nucleotide diversity is associated with a high likelihood of mutation and codons under diversifying selection. **A**. Average nonsynonymous pairwise nucleotide diversity per site (*π*_N_) and average synonymous pairwise nucleotide diversity per site (*π*_S_). **B**. *π*_S_ vs *π*_N_ of the coding sequence of NLRs with per group linear regressions. **C**. Mutation probability score of hv and non-hvNLRs. **D**. Percentage of codons under positive selection determined by MEME (episodic), and FEL (pervasive).

Since elevated *π*_N_ and *π*_S_ with no difference in *π*_N_/*π*_S_ could be caused by an increase in mutation rate of hvNLRs, we compared the predicted SNVs and indels per base pair based on epigenomic states (mutation probability score) (Monroe *et al*., 2022). The mutation probability is 35% higher for hvNLRs (**Fig 5C**; unpaired wilcoxon rank sum test, p=3.0e-05).

Gene-wide *π*_N_/*π*_S_ is a conservative metric for testing positive selection because positive selection may only be acting at a few codon sites (Kosakovsky Pond and Frost, 2005). Therefore, we used maximum-likelihood based site models to test for positive, diversifying selection. Use of these dN/dS-based models on intraspecies data is problematic because the nucleotide differences do not represent substitutions fixed by selection, but rather polymorphisms segregating within a population (Kryazhimskiy and Plotkin, 2008). We mitigated this effect by restricting our analysis to internal branches of the protein phylogeny, which encompass at least one ancestral sequence that is visible to selection (Pond *et al*., 2006; Avanzato *et al*., 2019). hvNLRs have a higher proportion of codons under pervasive and episodic diversifying selection than non-hvNLRs, indicating that diversifying selection at a subset of sites is maintaining diversity at hvNLRs (**Fig. 5D**, unpaired wilcoxon rank sum test). Given the polymorphism data, summary statistics, and mutational likelihood, hvNLR amino acid diversity appears to be driven by both a higher likelihood of mutation and positive, diversifying selection, while non-hvNLR conservation is maintained by purifying selection.

As described previously, hv and non-hvNLRs can co-exist as neighboring genes. We chose non-hvNLR *AT5G43730* and hvNLR *AT5G43740*, two CNLs of similar length 1.8kb apart, to examine the genomic features and signatures of selection of neighboring NLRs (**Fig. 6**). The hvNLR is highly expressed, hypomethylated, and has a TE within its 5’ UTR sequence (**Fig. 6A**). The non-hvNLR shows signatures of purifying selection with a gene-wide Tajima’s D value of -1.9, while the hvNLR has a gene-wide Tajima’s D of -0.24 (**Fig. 6B, 6C**). The hvNLR has higher *π, π*_N_, and *π*_S_, but the two genes have similar *π*_N_/*π*_S_ values (0.48 and 0.41) (**Fig. 6B, 6C**). Despite neighboring genomic positions, *AT5G43730* and *AT5G43740* show distinct genomic features and signatures of selection reflective of their species-level amino acid diversity (**Fig. 6B, 6C**). Therefore, we conclude that genomic features that distinguish hvNLR and non-hvNLRs are not driven by broader genome states, but may instead be related to function and evolutionary speed.

**Figure 6:**
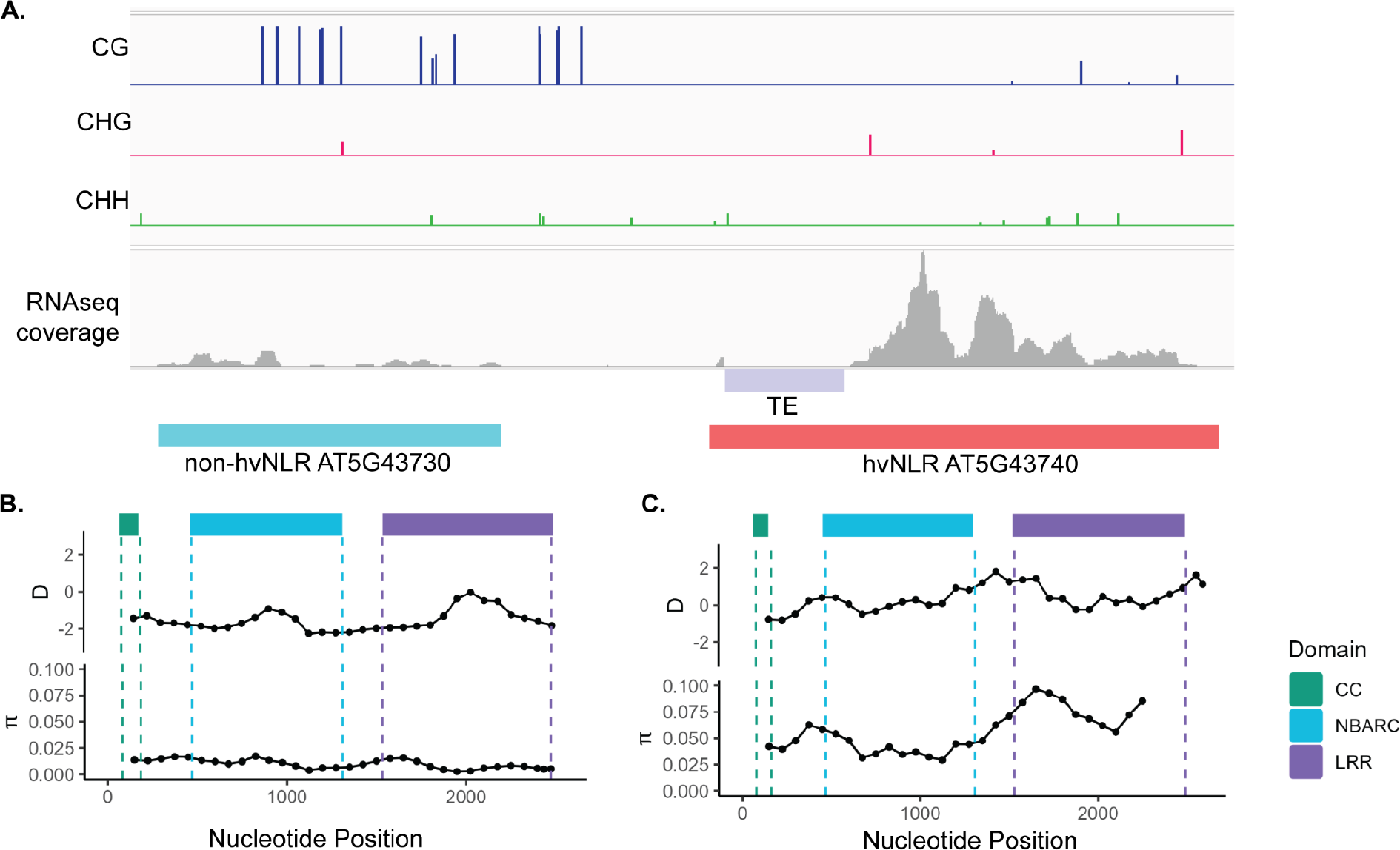
Neighboring NLRs retain distinct genomic and epigenomic features. **A**. Methylation, RNAseq coverage, and TE proximity of neighboring non-hvNLR *AT5G43730* and hvNLR *AT5G43740*. **B**. and **C**. Tajima’s D and nucleotide diversity across the coding sequence of *AT5G43730* and *AT5G43740*. Statistics were calculated on 300bp windows with a step size of 75bp, and plotted at the nucleotide midpoint.

## Discussion

The high allelic diversity of NLRs has long been appreciated, though the mechanisms that generate and maintain this diversity have remained difficult to disentangle. Taking advantage of Shannon entropy and available long read sequencing datasets, we can delineate rapidly and slowly diversifying NLRs and begin to investigate these mechanisms through gene set comparison. Our results show that rapidly evolving NLRs have distinct genomic features from their conserved paralogs and the rest of the genome. Specifically, we found that hvNLRs are more expressed, less methylated, and closer to TEs than non-hvNLRs. Interestingly, hvNLRs are enriched across the genome in highly expressed genes, hypomethylated genes, and genes closest to TEs, while non-hvNLRs are uniformly dispersed among other genes.

Since we observed distinct genomic features between hv and non-hvNLRs, we investigated the possibility of increased mutation rate in hvNLRs through examination of nucleotide diversity and mutation probability. Synonymous substitutions are under reduced selection compared to nonsynonymous substitutions because they do not alter the amino acid sequence, but are not invisible to selection due to codon bias, GC biased gene conversion, and RNA folding stability (Martincorena, Seshasayee and Luscombe, 2012; James, Castellano and Eyre-Walker, 2017; Wei, 2020). *π*_S_ is therefore an imperfect predictor of mutation rate, but an elevated mutation rate of hvNLRs could result in increased *π*_S_ and *π*_N_ relative to non-hvNLRs, but not influence the *π*_N_/*π*_S_ ratio, as we report here (Bromham, Cowman and Lanfear, 2013). We also find that hvNLRs are maintained in chromatin states associated with a higher mutation probability per base pair relative to non-hvNLRs, leading to the hypothesis that locally high mutation rate at hvNLRs contributes to the observed amino acid diversity. However, high depth quantification of *de novo* mutations at NLRs before selection is required to evaluate this hypothesis.

The distinct genomic features between the two NLR groups may point to mechanisms of increased mutation rate. Transcription is a source of genomic instability through the exposure of vulnerable single-stranded DNA, which is countered by targeting DNA repair machinery to actively transcribed genes through the stalling RNA polymerase or histone marks associated with actively transcribed genes (Oztas *et al*., 2018; Quiroz *et al*., 2022). If the high transcription of hvNLRs is not accompanied by targeted DNA repair, this would result in an increased probability of mutation (Staunton, Peters and Seoighe, 2023). Methylated cytosines increase the likelihood of mutation by increasing the frequency of spontaneous deamination of cytosines (Xia, Han and Zhao, 2012; Weng *et al*., 2019; Monroe *et al*., 2022). However, in *Arabidopsis*, gene body CG methylation is found preferentially in the exons of conserved, constitutively transcribed housekeeping genes, and gene body CG methylation is associated with lower polymorphism than unmethylated genes across accessions (Gaut *et al*., 2011; He *et al*., 2022; Kenchanmane Raju *et al*., 2023). The CG gene body methylation of non-hvNLRs may therefore be related to their low diversity through some unknown mechanism. TEs generate large effect mutations (Quadrana *et al*., 2019) and alter the methylation and expression landscape of surrounding genes. hvNLRs are closer to TEs and more likely to have them within their genic sequence than non-hvNLRs, and this likely contributes to hvNLR diversification.

Once generated, nucleotide diversity can be actively maintained by diversifying or balancing selection, or passively accumulate in the absence of selection. We do not observe any difference in diversifying selection between hv and non-hvNLRs using the *π*_N_/*π*_S_ metric, but hvNLRs have a significantly higher proportion of codons under pervasive and episodic diversifying selection. While hvNLRs have higher Tajima’s D values than the genome average and non-hvNLRs, they are not present in the tails of the genome-wide distribution. The 5th and 95th percentiles of the empirical distribution is a conservative cutoff, and it is possible for a locus to be under selection but not in a tail of the distribution if selection is weak. Therefore, balancing selection may play a role in promoting hvNLR diversity, but cannot be distinguished from evolution under relaxed selection using this criteria. non-hvNLRs, however, have a strong signature of purifying selection, which helps to explain their low amino acid diversity relative to hvNLRs.

Given the heterogeneous mutation rate across the *Arabidopsis* genome, it is tempting to speculate that the distinctive genomic features we observed in hvNLRs may be related to their allelic diversity. Alternatively, there might be a selection of specific features on non-hvNLRs to enhance DNA repair and inhibit other diversity-generation activities facilitating their maintenance. Our findings serve as a starting point for the investigation of the mechanisms that promote diversity generation in a subset of the plant immune receptors.

## Materials and Methods

To examine the methylation and expression of NLRs, we used available matched bisulfite and RNA sequencing from split Col-0 leaves (Williams *et al*., 2022). Reads were trimmed using Trim Galore! v0.6.6 with a Phred score cutoff of 20 and Illumina adapter sequences, with a maximum trimming error rate 0.1 (Babraham Bioinformatics). Using Bismark v0.23.0, reads were mapped to the Araport11 genome, PCR duplicates were removed, and percent methylation at each cytosine was determined using the methylation extraction function (Krueger and Andrews, 2011). Cytosines with at least 5 reads were used for analysis, and the symmetrical cytosines within CG base pairs were averaged (Williams *et al*., 2022). The percent methylation of each CG site was averaged across each NLR gene, and across four biological replicates. Five hvNLR genes did not have sufficient coverage at any cytosines and were excluded from analysis *(AT1G58807, AT1G58848, AT1G59124, AT1G59218*, and *AT4G26090*).

RNA-seq reads from four matched leaf samples (explained above) were mapped to the Araport11 genome using STAR v2.7.10a and were counted using htseq-count v2.0.2 (Dobin *et al*., 2013). Counts were converted to transcripts per million and averaged across four biological replicates, then log2 transformed for visualization. NLRs are repetitive and often similar, making them difficult to sequence with short reads. To determine if any NLRs were unmappable, RNAseq reads were simulated using Polyester v1.2.0 (Frazee *et al*., 2015). Four NLRs were determined to be unmappable due to zero assigned read counts and were excluded from expression analysis (*AT1G58807, AT1G58848, AT1G59124*, and *AT1G59218*). Single sample gene set enrichment of hvNLRs and non-hvNLRs was performed on each replicate using singscore (Foroutan *et al*., 2018).

We determined distance to transposable elements based on the TE annotation file TAIR10_Transposable_Elements.txt and gene annotation file TAIR10_GFF3_genes.gff available from arabidopsis.org. The phylogenetic tree of all NLRs in Col-0 was generated as described previously (Prigozhin and Krasileva, 2021) with feature annotations using iTOL. The UpSet plot was generated using the R package ComplexUpset v1.3.3.

Protein alignments for each NLR were generated as described previously (Prigozhin and Krasileva, 2021) and converted to codon alignments using PAL2NAL v14 (Suyama, Torrents and Bork, 2006). The population genetics statistics of NLRs were calculated using EggLib v3.1.0 (Siol *et al*., 2022). Domain specific statistics were calculated on subsets of codon alignments using majority vote across annotations. NB-ARC, TIR, and CC annotations were collected from previous work (Van de Weyer *et al*., 2019), and LRR annotations were determined using LRRpredictor (Martin *et al*., 2020). Sliding window analysis was performed with 300 base pair windows with a 75 base pair step. Sites under pervasive diversifying selection were identified using FEL (Kosakovsky Pond and Frost, 2005) and sites under episodic diversifying selection were identified using MEME (Murrell *et al*., 2012) using the internal branches of the phylogeny. Empirical distributions of population genetics statistics of coding sequences were calculated from the all sites 1001 Genomes VCF subset to the accessions used to generate the NLRome long read dataset using vcftools v0.1.17 (Danecek *et al*., 2011; Alonso-Blanco *et al*., 2016; Van de Weyer *et al*., 2019).

All the data generated in this study is hosted on the Zenodo Public Repository at 10.5281/zenodo.7527904. The processing pipelines and figure generation code are available on Github (https://github.com/chandlersutherland/nlr_features).

## Acknowledgements

We are grateful to the Krasileva Lab for the critical reading of the manuscript. This research used the Savio computational cluster resource provided by the Berkeley Research Computing program at the University of California, Berkeley (supported by the UC Berkeley Chancellor, Vice Chancellor for Research, and Chief Information Officer). Chandler A. Sutherland has been supported by the Grace Kase-Tsujimoto Graduate Fellowship. Ksenia V Krasileva is funded by NIH Director’s Award (1DP2AT011967-01), Gordon and Betty Moore Inventor Fellowship (grant number: 8802) and the Innovative Genomics Institute.

## Supplemental Figures

**Supplemental Figure 1:**
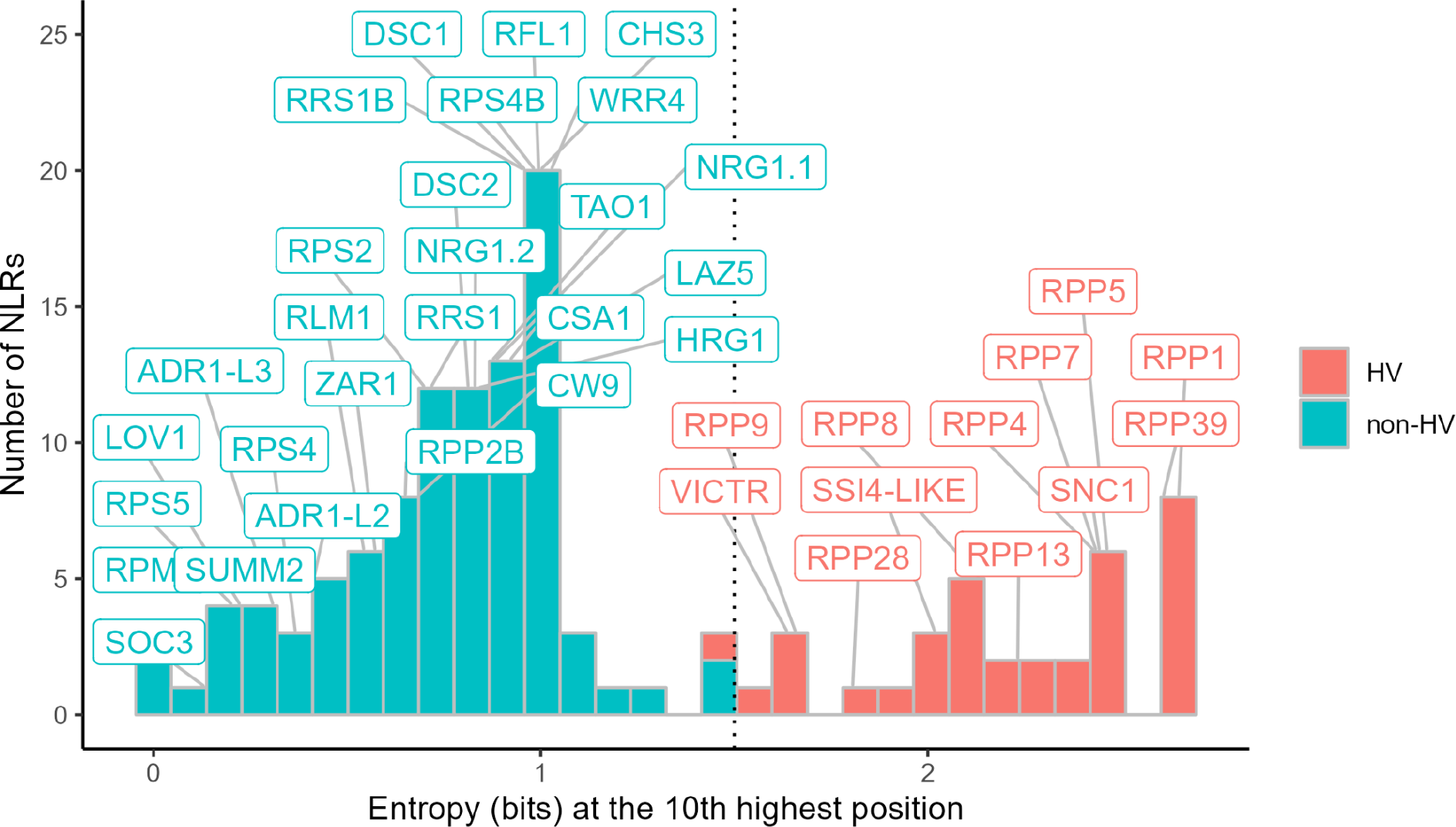
Distribution of NLR Shannon entropy at the tenth highest amino acid position as shown as a histogram with 30 bins. Described NLRs are annotated. The designation of hvNLR is entropy of >1.5 bits at the tenth highest position, as shown by the dashed line.

**Supplemental Figure 2:**
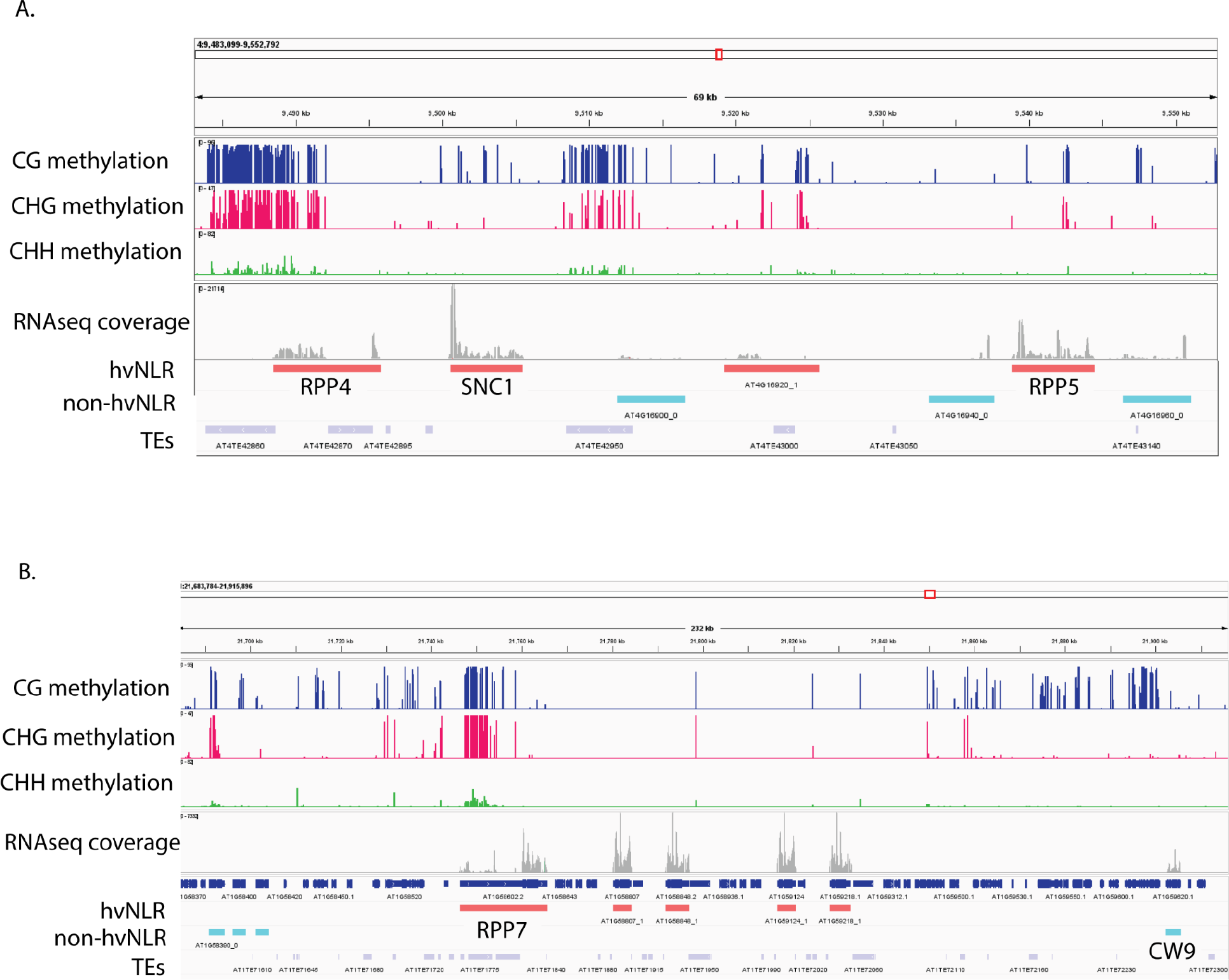
Integrative Genomics Viewer screenshot of Methylation, RNAseq coverage, and TE proximity of the **A**. *RPP4* and **B**. *RPP7* clusters.

**Supplemental Figure 3:**
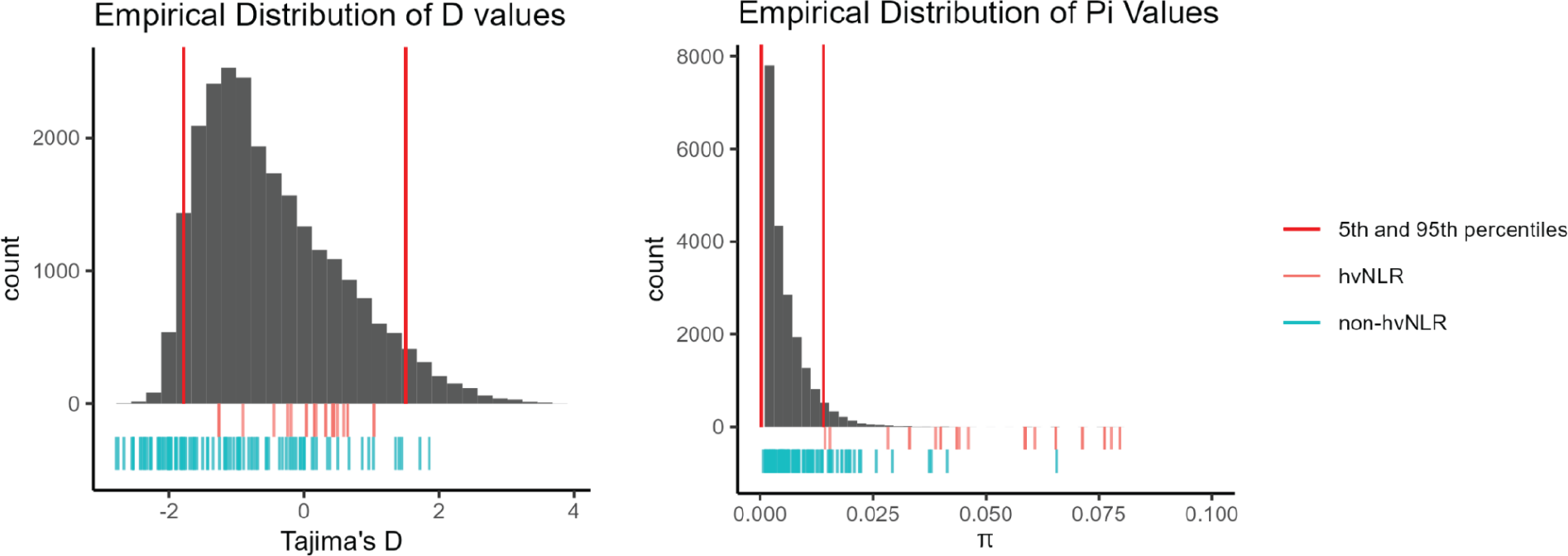
Empirical distribution of Tajima’s D and *π* calculated on coding sequences of *Arabidopsis*. Position of hv and non-hvNLRs shown via rug plot, as well as the 5th and 95th percentiles of the distribution.

**Supplemental Figure 4:**
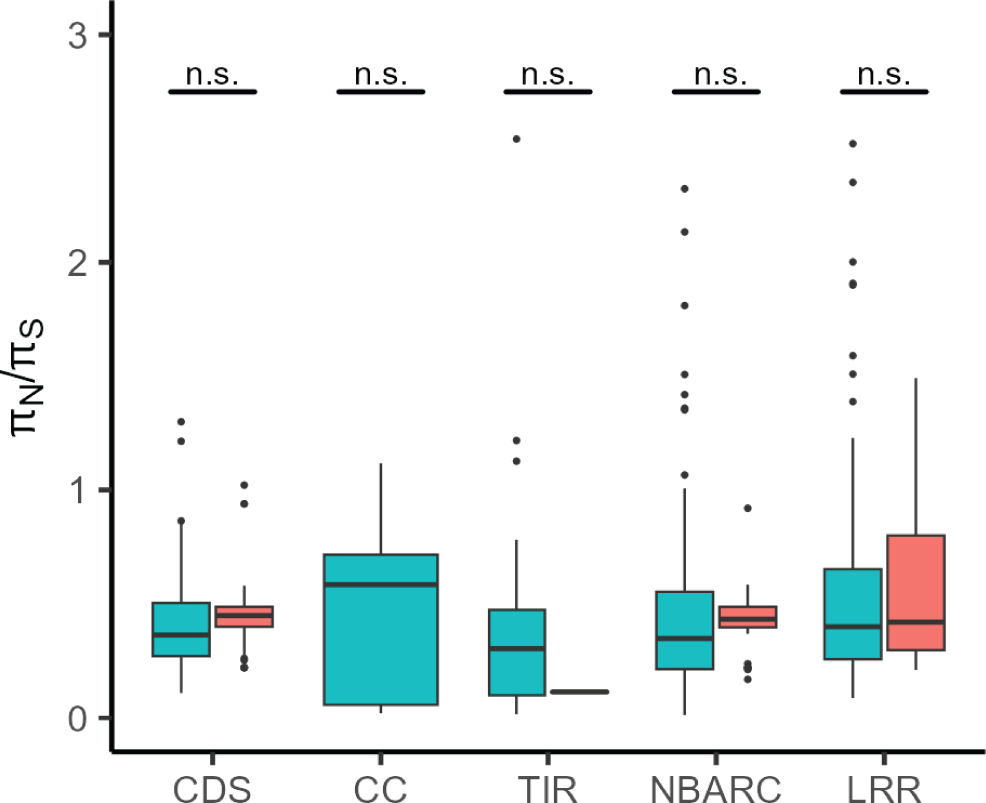
*π*_N_/*π*_S_ calculated per domain.

